# Highly multiplexed immunofluorescence images and single-cell data of immune markers in tonsil and lung cancer

**DOI:** 10.1101/704114

**Authors:** Rumana Rashid, Giorgio Gaglia, Yu-An Chen, Jia-Ren Lin, Ziming Du, Zoltan Maliga, Denis Schapiro, Clarence Yapp, Jeremy Muhlich, Artem Sokolov, Peter Sorger, Sandro Santagata

**Affiliations:** Department of Pathology, Brigham and Women’s Hospital, Harvard Medical School, Boston, MA; Laboratory for Systems Pharmacology, Harvard Medical School, Boston, MA; Ludwig Center at Harvard, Harvard Medical School, Boston, MA; Department of Biomedical Informatics, Harvard Medical School, Boston, MA; Broad Institute of MIT and Harvard, Cambridge, MA; Department of Systems Biology, Harvard Medical School, Boston, MA; Department of Oncologic Pathology, Dana Farber Cancer Institute, Boston, MA

**Keywords:** multiplexed imaging, tissue imaging, quantitative pathology, single-cell analysis, image analysis, immunotherapy, antibody qualification, formaldehyde-fixed paraffin-embedded, FFPE, immune profiling, immune checkpoint, PD-1, PD-L1, antibody, immunofluorescence, tissue based cyclic immunofluorescence, t-CyCIF, spatial phenotyping, FOXP3

## Abstract

In this data descriptor, we document a dataset of multiplexed immunofluorescence images and derived single-cell measurements of immune lineage and other markers in formaldehyde-fixed and paraffin-embedded (FFPE) human tonsil and lung cancer tissue. We used tissue cyclic immunofluorescence (t-CyCIF) to generate fluorescence images which we artifact corrected using the BaSiC tool, stitched and registered using the ASHLAR algorithm, and segmented using ilastik software and MATLAB. We extracted single-cell features from these images using HistoCAT software. The resulting dataset can be visualized using image browsers and analyzed using high-dimensional, single-cell methods. This dataset is a valuable resource for biological discovery of the immune system in normal and diseased states as well as for the development of multiplexed image analysis and viewing tools.

**METADATA SUMMARY:** 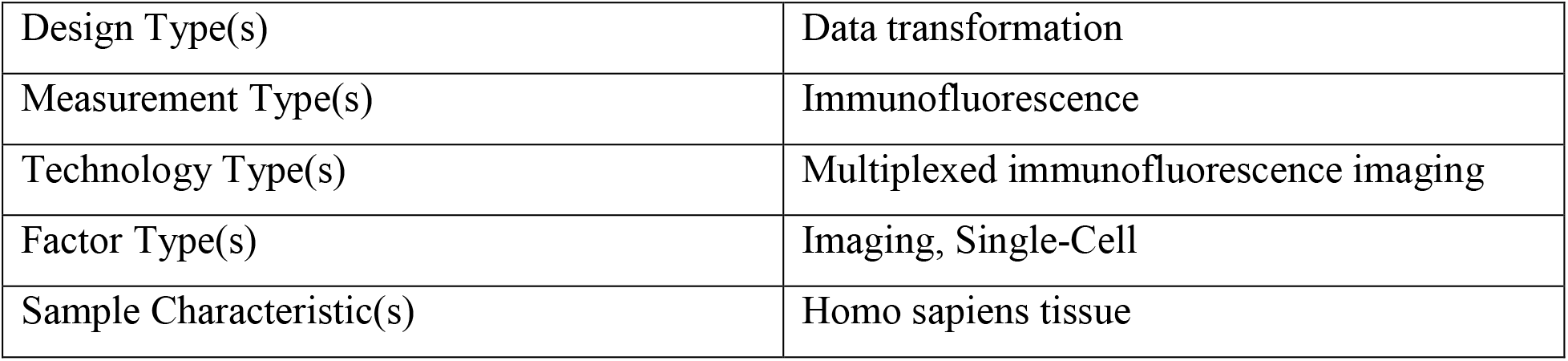

## BACKGROUND & SUMMARY

Tissues comprise individual cells of diverse types along with supportive membranes and structures as well as blood and lymphatic vessels. The identities, properties and spatial distributions of cells that make up tissues are still not fully known: classical histology provides excellent spatial resolution, but it typically lacks molecular details. As a result, the impact of intrinsic factors such as lineage and extrinsic factors such as the microenvironment on tissue biology in health and disease requires molecular profiling of single cells within the broader context of organized tissue architecture. Such deep spatial and molecular phenotyping is especially pertinent to the study of cancer resection tissues. These samples are routinely acquired prior to, on, and after a therapeutic intervention, providing opportunities to characterize the interplay between malignant tumor cells and surrounding immune cell populations and how those relationships are influenced over time by treatments. Understanding these relationships may provide biomarker signatures that predict response to therapy^1,2^ and is particularly relevant in the case of immunotherapeutics. Many available immunotherapies, including those targeting cytotoxic T lymphocyte-associated antigen-4 (CTLA-4), programmed cell death-1 receptor (PD-1), and programmed cell death-1 ligand (PD-L1), influence interactions between tumor and immune cells to inhibit immune checkpoints and activate the immune system’s surveillance of tumor cells^3–7^. However, even in tumor types that are highly responsive to such therapies, many patients do not benefit, and many types of tumors remain broadly refractory to these agents. A deeper understanding of immune cell states, location, interactions, and architecture (“immunophenotypes”) promises to provide new prognostic and predictive information for cancer research and treatment.

With recent advances in multiplexed imaging technologies^8^, multiple epitopes can be detected within a tissue section and the spatial distributions and interactions of cell populations precisely mapped. One such method is tissue-based cyclic immunofluorescence (t-CyCIF)^9^ which yields high-plex images at subcellular resolution and has been used to characterize immune populations in several tumor types^10–12^. In t-CyCIF, a high-plex image is constructed from a series of 4 to 6 color images, which are then registered and superimposed. The images provide information on the amount of an epitope that is expressed as well as the location of the epitope within the tissue. By segmenting the images to demarcate single cells or subcellular compartments, we can then use epitope expression levels to discriminate immune, tumor, and stromal cell types and compute their numbers and distributions within tumors and surrounding normal tissue.

The quality of the antibody reagents largely dictates the reliability of data that is generated by antibody-based imaging methods such as multiplexed ion beam imaging (MIBI)^13^, imaging mass cytometry (IMC)^14^, co-detection by indexing (CODEX)^15^, DNA exchange imaging (DEI)^16^, MultiOmyx (MxIF)^17^, imaging cycler microscopy (ICM)^18–20^, multiplexed IHC^21^, NanoString Digital Spatial Profiling (DSP)^22^, and t-CyCIF itself. We have recently published detailed methods for validating antibodies and assembling panels of antibodies for multiplexed tissue techniques in Du, Lin, Rashid *et al*., (Nature Protocols 2019, in press)^23^. That work highlights a variety of complementary approaches to qualify antibodies using information at the level of pixels, cells, and tissues and yielded a 16-plex antibody panel capable of detecting lymphocytes, macrophages, and immune checkpoint regulators for use in ‘immune profiling’ tissue sections. Using t-CyCIF, we qualified antibodies in reactive tonsil tissue (TONSIL-1), which has a highly stereotyped arrangements of diverse immune cell types, and then demonstrated the panel’s utility in characterizing common and rare immune populations in three lung cancer tissue specimens: a lung adenocarcinoma that had metastasized to a lymph node (LUNG-1-LN), a lung squamous cell carcinoma that had metastasized to the brain (LUNG-2-BR), and a primary lung squamous cell carcinoma (LUNG-3-PR). We also provide t-CyCIF imaging data from 8 FFPE sections used to validate antibodies; in these samples, antibodies were used in different permutations and order, making the data useful for examining relationships between antigenicity, fluorescence signal, and cycle number.

In this data descriptor, we share the images from Du, Lin, Rashid *et al*., (Nature Protocols 2019, in press)^23^. The dataset includes immunofluorescence images from formalin fixed paraffin embedded (FFPE) tissue sections mounted onto glass slides. In each section, there are between ~61,800 to ~483,000 individual cells with fluorescence intensity and spatial information provided for 27 antibodies that were acquired in a multiplexed fashion. These antibodies include the highly validated 16-plex panel as well as antibodies against several additional markers of interest such as markers of tumor cell lineage and cell proliferation. We also include quantitative, single-cell measurements of 60+ features including fluorescence intensity measurements for each target epitope/protein, cellular morphology measurements such as area, eccentricity, and solidity, and spatial information such as the centroid position of a cell and its nearest neighbors.

The resulting single-cell data can be analyzed using quantitative and qualitative approaches both in the context of the original spatial arrangement of the tissue and as sets of derived feature vectors, one for each cell. Spatial views enable the analysis of geographic patterns and interactions between different cells types, such as the immune microenvironment surrounding tumor tissue. Such data can be used to develop new methods for visualizing large complex images and to develop and refine data analysis approaches such as image segmentation, intensity gating (to discriminate ‘positive’ and ‘negative’ cell populations), and spatial clustering. As multiple research centers begin to assemble high-dimensional and multi-parametric atlases of human cancers and pre-cancers, there is an increasing need for cross-center validation of analysis methodologies. Publicly available datasets such as ours will provide a freely accessible resource for such efforts.

## METHODS

### Tissue Samples

Five formalin-fixed paraffin-embedded (FFPE) human tissue samples were retrieved from the archives of the Department of Pathology at Brigham and Women’s Hospital with IRB approval as part of a discarded tissue protocol. The diagnoses were confirmed by a board-certified pathologist (S.S.) (Table 1). Sections were cut from FFPE blocks at a thickness of 5 μm and mounted onto Superfrost Plus microscope slides prior to use.

### Datasets

Data from tissue samples was acquired in two batches. The first batch (DATASET-1) contains data from LUNG-1-LN, LUNG-2-BR, LUNG-3-PR, and TONSIL-1. The second batch (DATASET-2) contains data from eight sections of TONSIL-2. Data associated with each of these sections are labeled TONSIL-2.1, TONSIL-2.2, etc. in the data records.

### Tissue-Based Cyclic Immunofluorescence

Each section of tissue was imaged with a panel of 26-28 antibodies using t-CyCIF as previously described^9^. This method consists of iterative cycles of antibody incubation, imaging, and fluorophore inactivation (Figure 1).

**Figure 1.**
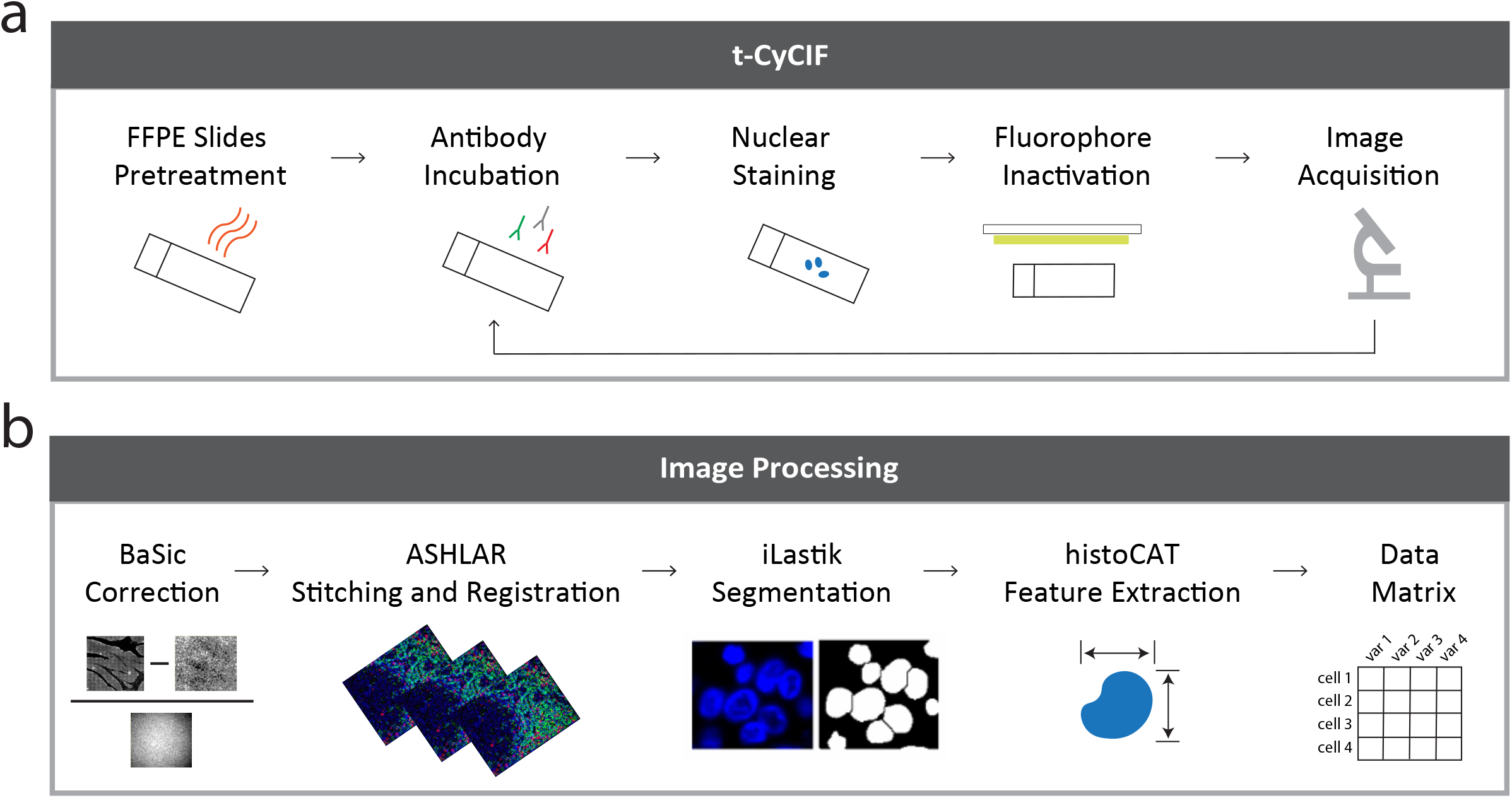
Overview of data generation. (a) Multiplexed, immunofluorescence images were acquired using the tissue-based cyclic immunofluorescence (t-CyCIF) method and (b) processed with a series of algorithms and toolboxes including BaSiC, ASHLAR, ilastik, and histoCAT to obtain single-cell features.

#### Slide Preparation

An automated program on the Leica Bond RX (Leica Biosystems) was used to prepare slides for t-CyCIF. The slides were treated as follows: baked at 60 °C for 30 min, dewaxed at 72 °C with Bond Dewax Solution (Cat. AR9222, Leica Biosystems), and treated with Epitope Retrieval 1 (ER1) Solution at 100 °C for 20 min for antigen retrieval. Odyssey Blocking Buffer (Cat. 927–40150, LI-COR) was applied to the slides at room temperature (RT) for 30 min and then incubated with three secondary antibodies at RT for 60 min, followed by Hoechst 33342 (Cat. H3570, Life Technologies) solution (2 ug/ml) at RT for 30 min.

#### Blocking

After slide preparation, non-specific, reactive epitopes were blocked by incubating slides overnight at 4 °C in the dark with secondary antibodies raised against the host species of the unconjugated, primary antibodies used in the first cycle of t-CyCIF.

#### Antibody Staining

Slides were initially imaged to measure nonspecific binding from secondary antibodies, photobleached, and then imaged again to measure autofluorescence. In the first cycle of antibody incubation, the slides were incubated overnight with primary antibodies from different species and then with secondary antibodies for two hours at RT in the dark. We washed slides with 1X PBS, stained them with Hoechst solution, and then imaged them. This process was repeated for 11-12 cycles using antibodies directly conjugated to fluorophores. All antibodies used in this study are listed in Table 2 with an assigned unique identifier. The antibodies used for each cycle of imaging for all samples in DATASET-1 are detailed in Table 3, and antibodies for all samples in DATASET-2 are detailed in Table 4.

#### Mounting and De-Coverslipping

Prior to each cycle of imaging, slides were wet-mounted using 200 μl of 10% glycerol in PBS and 24 x 50mm glass cover slips (Cat # 48393-081, VWR). Following imaging, slides were decoverslipped by placing vertically in a slide rack completely submerged in a container of 1X PBS for 15 minutes and slowly pulling the slides back up, allowing the glass coverslip to remain in the PBS.

#### Image Acquisition

Images from each cycle of t-CyCIF were acquired using the RareCyte CyteFinder Slide Scanning Fluorescence Microscope. The four following filter sets were used: 1) The ‘DAPI channel’ for imaging Hoechst with a peak excitation of 390 nm and half-width of 18 nm and a peak emission of 435 nm and half-width of 48 nm, 2) the ‘488 channel’ with a 475-nm/28-nm excitation filter and a 525-nm/48-nm emission filter, 3) the ‘555 channel’ with a 542-nm/27-nm excitation filter and a 597-nm/45-nm emission filter, and 4) the ‘647 channel’ with a 632-nm/22-nm excitation filter and a 679-nm/34-nm emission filter. Each tissue section was imaged twice, a large region with a 10X/0.3 NA objective and a smaller region with a 40X/0.6NA objective. The 10X images have a field of view of 1.6 x 1.4 mm and a nominal resolution of 1.06 μm. The 40X images have a field of view of 0.42 x 0.35 mm and a nominal resolution of 0.53 μm. For both images, a 5% overlap was collected between fields of view to facilitate image stitching. In DATASET-2, the first cycle of antibodies were imaged twice, once with a high exposure time and once with a low exposure time.

#### Photobleaching

Following slide preparation using the Leica Bond RX and subsequent to each cycle of imaging, fluorophores were inactivated by submerging slides in a solution of 4.5% H_2_O_2_ and 20mM NaOH in 1X PBS and incubating them under a light emitting diode (LED) for 2 hours at RT.

### Image Processing

#### Background and Shading Correction

The BaSiC algorithm^24^ plugin for ImageJ was used to computationally derive flat-field and dark-field profiles from the original image for each cycle. The flat-field is used to correct for irregular illumination of the sample, and the dark-field is used to correct for camera sensor offset and internal noise. Lambda values of 0.1 and 0.01were used for flat-field and dark-field, respectively. For each cycle, the raw image was subtracted by the dark-field profile and divided by the flat-field profile to correct the shading on each individual image field.

#### Stitching and Registration

Ashlar (version v1.6.0) was used to stitch the fields from the first imaging cycle into a mosaic and co-register the fields from successive cycles of imaging. Ashlar stitches fields together by calculating the phase correlation between neighboring images to correct for local state positioning error, and applying a statistical model of microscope stage behavior to correct for large-scale error. It uses a similar phase correlation approach to then register fileds from successive cycles to the first cycle of stitched images. The output is an OME-TIFF file that contains a seamless many-channel mosaic depicting the entire sample across all image cycles.

#### Segmentation

The OME-TIFF output from ASHLAR was used to segment single cells in the images using the ilastik software program^25^ and MATLAB(version 2018a). The OME-TIFF was cropped into 6000 x 6000 pixel regions to increase processing speed. From each cropped region, ~20 random 250 x 250 pixel regions were selected and used as training data on the ilastik program to generate a probability of each pixel in the cropped region belonging to three classes: nuclear area, cytoplasmic area, or area not occupied by a cell (background). Color/intensity features including gaussian smoothing, edge features including the Laplacian of gaussian, gaussian of gradient magnitude, and difference of gaussians, and texture features including structure tensor eigenvalues and hessian of gaussian eigenvalues with a σ_0_ = 0.30, σ_1_ = 0.70, σ_2_ = 1.00, σ_3_ = 1.60, σ_4_ = 3.50, and σ_5_ = 5.03 were used for pixel classification. MATLAB was then used to perform a watershed segmentation on the probabilities to identify objects, or cells, and a segmentation mask was generated for each cropped region.

#### Single-Cell Feature Extraction

The histology topography cytometry analysis toolbox (histoCAT)^26^ was used to extract features of the cells segmented in each image. Single cell features included fluorescence intensity measurements of each antibody, morphological features such as cell area and circularity, as well as spatial features such as the centroid position of the cell. Moreover, cells in spatial proximity to one another were identified and indexed to enable neighborhood analysis and cell phenotype interactions. The output was a data table for each cropped region. For each sample, the data tables from all the cropped regions were concatenated into a master image level data table with each cell assigned a global unique identifier and centroid position. A complete list and description of each feature in the master data tables is provided in Table 5.

### Code availability

All code used to process and generate the data in this study can be found alongside the data (Data Citation 1: Synapse https://doi.org/10.7303/syn17865732)^27^. Source code for ASHLAR is available on GitHub (https://github.com/jmuhlich/ashlar). The newest histoCAT version can also be found GitHub (https://github.com/BodenmillerGroup/histoCAT).

## DATA RECORDS

We have made all the data for this manuscript available in the Synapse repository hosted by Sage Bionetworks (Data Citation 1: Synapse https://doi.org/10.7303/syn17865732)^27^. We organized the data as described in Figure 2. For each tissue sample, we share image data acquired at two magnifications. For each 40X magnification dataset, we share:

i. raw rcpnl files,
ii. illumination profiles calculated by the BaSiC algorithm,
iii. an OME-TIFF file output from the ASHLAR algorithm,
iv. individual TIFF images for each marker,
v. probability masks for segmentation from ilastik software,
vi. labeled nuclear segmentation mask, and
vii. data table of 60+ features extracted for each cell.

**Figure 2.**
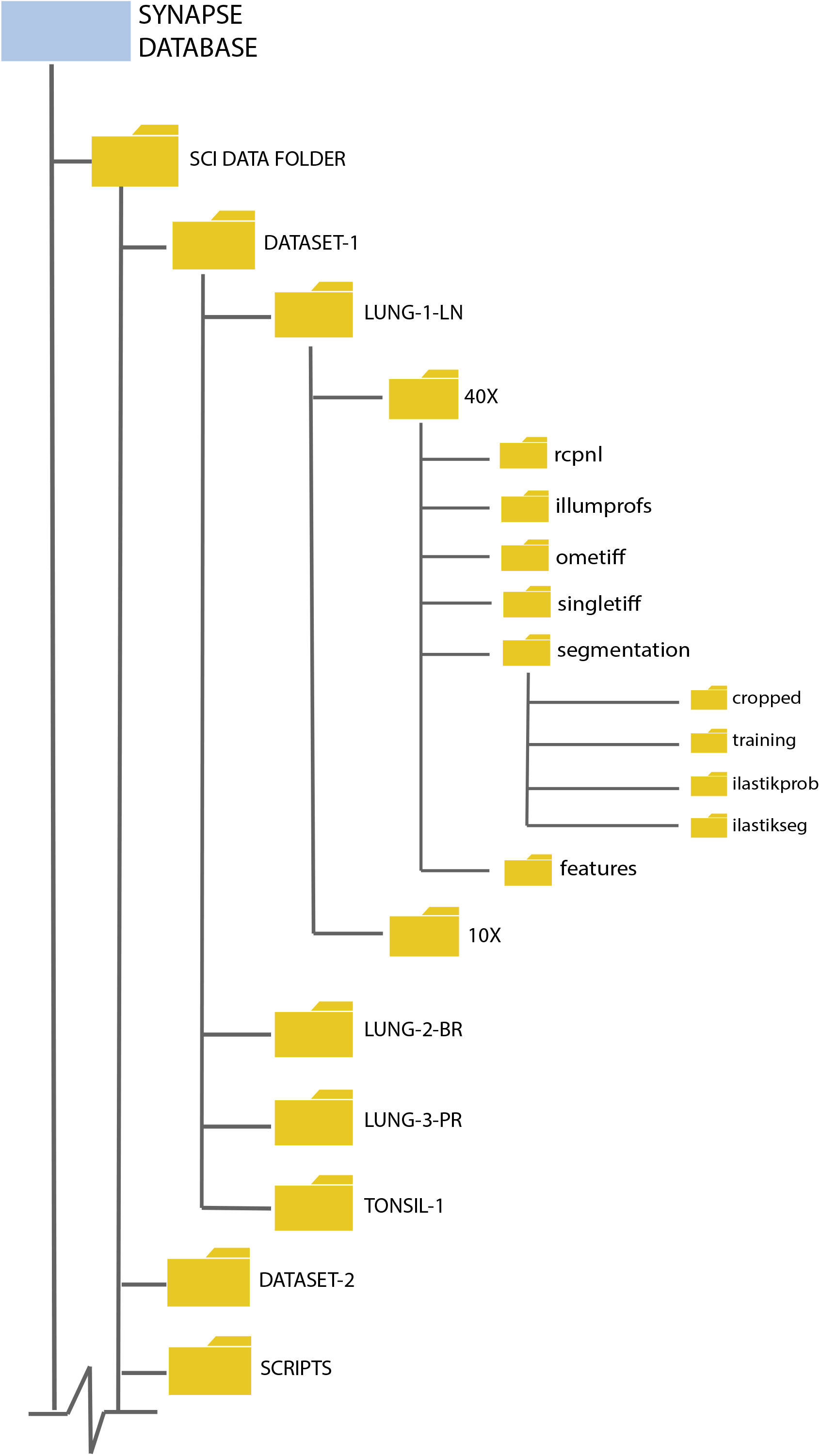
Database structure. All shared data are stored in the SYNAPSE repository. https://doi.org/10.7303/syn17865732

The “rcpnl” folder contains the raw image files in an rcpnl file format generated by the RareCyte CyteFinder for each cycle of imaging. The “illumprofs” folder contains TIFF files for the dark-field profile and the flat-field profile for each cycle of imaging. Each TIFF file in this folder is a stack of four TIFF images corresponding to the four wavelengths imaged every cycle. The “ometiff ‘ folder contains one OME-TIFF file that is a stitched, registered mosaic of all channels from all cycles of imaging. The OME-TIFF file contains mosaics at multiple resolutions. The “singletiff” folder contains a single TIFF mosaic for each marker. This folder separates the OME-TIFF into separate channels to facilitate opening in older software that is incompatible with the OME-TIFF format. The “segmentation” folder contains subfolders with intermediate data outputs from the segmentation process. The “cropped” subfolder contains 6000 x 6000 pixel regions from the OME-TIFF file. The “training” subfolder contains 250 x 250 pixel regions used as training data for segmentation. The “ilastikprob” subfolder contains a TIFF image for the probability of each pixel in the cropped regions belonging to each class used in ilastik training. This image is in a stack with the DAPI image from the first cycle of imaging for easy comparison of the accuracy of the probability mask. The “ilastikseg” folder contains a TIFF image of the nuclear segmentation mask. Each object, or cell, in the mask is labeled with a unique index number. The “features” folder contains a csv data table for each cropped region with 60+ feature measurements for each cell as well as a master data table with data from each cropped region combined.

We provide all scripts used in data generation. A description of the scripts and supporting documents is provided in Table 6.

Additionally, a subset of the imaging data can be found and viewed on cycif.org (Data Citation 2: CyCIF.org https://www.cycif.org/featured-paper/du-and-lin-2019/figures/). In this interactive image browser, we indicate some distinct regions of the tonsil and lung cancer images and provide descriptive narrations about a subset of the combinations of immune markers expressed in these samples.

## TECHNICAL VALIDATION

### Staining Quality

We performed a detailed validation of the panel of antibodies used to generate the datasets described in Du, Lin, Rashid *et al*., 2019^23^ One or more trained pathologists visually reviewed the staining patterns for each antibody to assess specificity to cell type, appropriate localization within the cell (nucleus v. cytoplasm v. membrane), co-staining with other markers, and localization to the expected geographic regions within the tissue. For example, the cytokeratin antibody, known to detect intermediate filament proteins in epithelial cells, was expressed in striated patterns surrounding the nuclei of cells morphologically consistent with epithelial origin, whereas the FOXP3 antibody, targeting a transcription factor in T cells, was concentrated in the nuclear area of small cells morphologically consistent with lymphocytes (Figure 3a). The antibodies detecting cell lineage markers such as FOXP3, which delineates a regulatory T-cell population, were further corroborated by assessing appropriate co-expression of other markers. For example, we found that FOXP3 was co-expressed with CD4, CD3D, and CD45, thereby increasing our confidence in the staining quality (Figure 3a). As another example, CD20, a B-cell antigen, was observed to have higher levels of signal within germinal centers of tonsil tissue which are well-established B cell rich compartments within tonsil rather than the mantle region where we found an abundance of cells expressing the T-cell antigen CD3D (Figure 3b). See Du, Lin, Rashid *et al*., 2019^23^ for additional quality measurements including the comparison of t-CyCIF antibody staining to the staining observed with clinical grade antibodies that were used in immunohistochemistry (IHC) staining, pixel-by-pixel correlations of multiple antibody clones against the same target, and various high-dimensional cell clustering methods.

**Figure 3.**
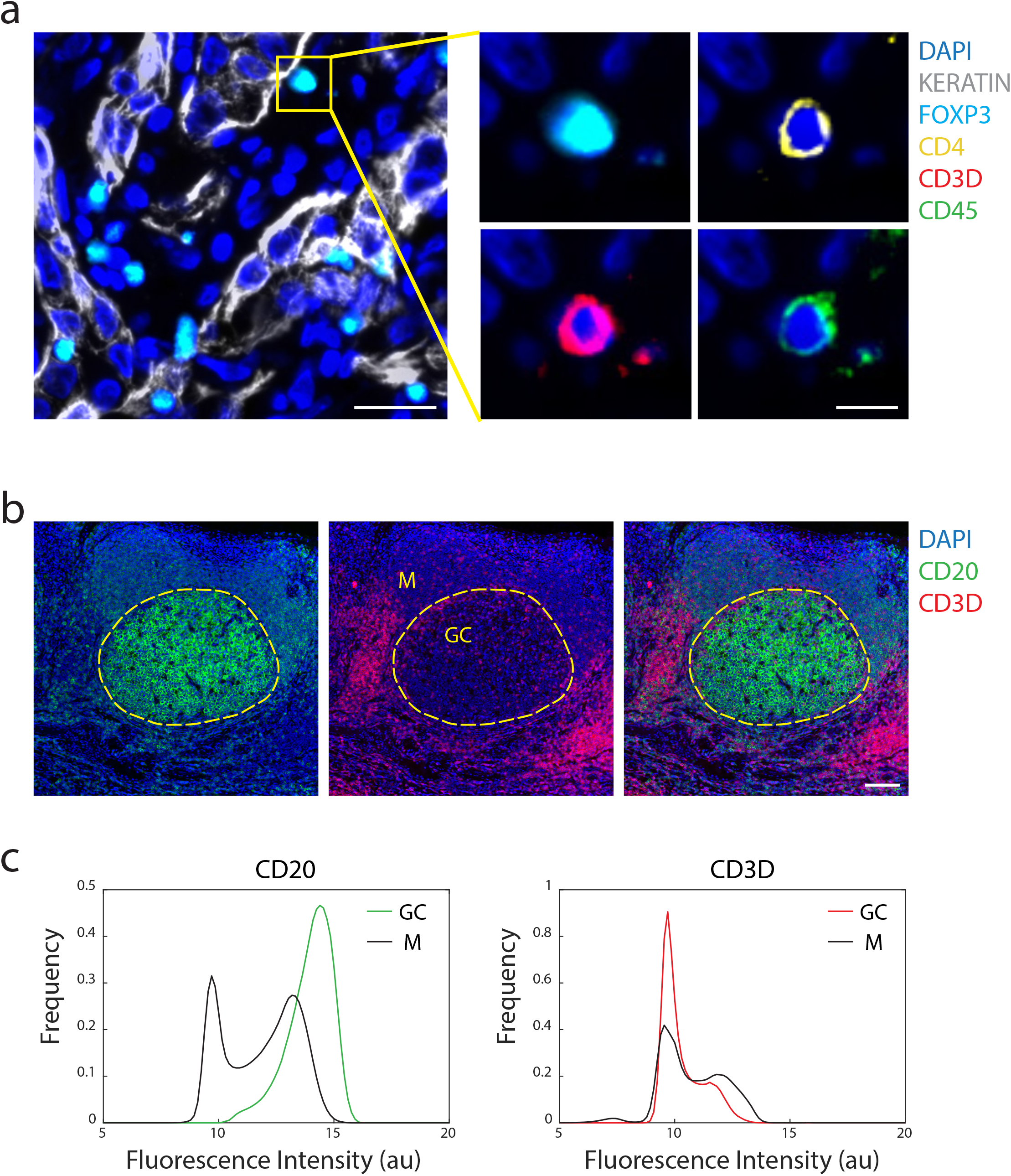
Antibody staining quality. (a) Immunofluorescence image from LUNG-3 showing epithelial tumor cells marked by Keratin (white) and a regulatory T cell marked by FOXP3 (cyan), CD4 (yellow), CD3D (red), and CD45 (green) (scale bar: 25 μm; inset scale bar: 10 μm). (b) A region of TONSIL-1 showing CD20 (green) and CD3D (red) expression. Area inside yellow dashed circle denotes germinal center (GC), and area outside denotes the mantle (M) region (scale bar: 100 μm). (c) Probability density function of fluorescence signal intensity of every pixel in the germinal center (n = 1,446,450 pixels) and mantle (n = 4,369,358 pixels) for CD20 and CD3D within the region shown in (a).

### Cell Segmentation

We evaluated the quality of segmentation of single cells within the tissue images using a two-step system. We only performed segmentation on the 40X magnification images because the lower resolution of the 10X magnification images did not give sufficient resolution. First, we overlaid the segmentation masks over the DAPI signal to evaluate the accuracy of segmentation qualitatively (Figure 4a); based on these data, we then adjusted and optimized the segmentation. Second, three users evaluated a random sample of 500 cells from the tonsil and each of the lung tissues to quantify the accuracy and rate of fusions (under-segmentation) and fission/splitting (over-segmentation) among mis-segmented cells (Figure 4b-c, Online-only Table 1). The cell segmentation of all samples had a low error rate (~0.1) across cells of various morphologies (large tumor cells, smaller round immune cells, skinny elongated fibroblasts, etc.).

**Figure 4.**
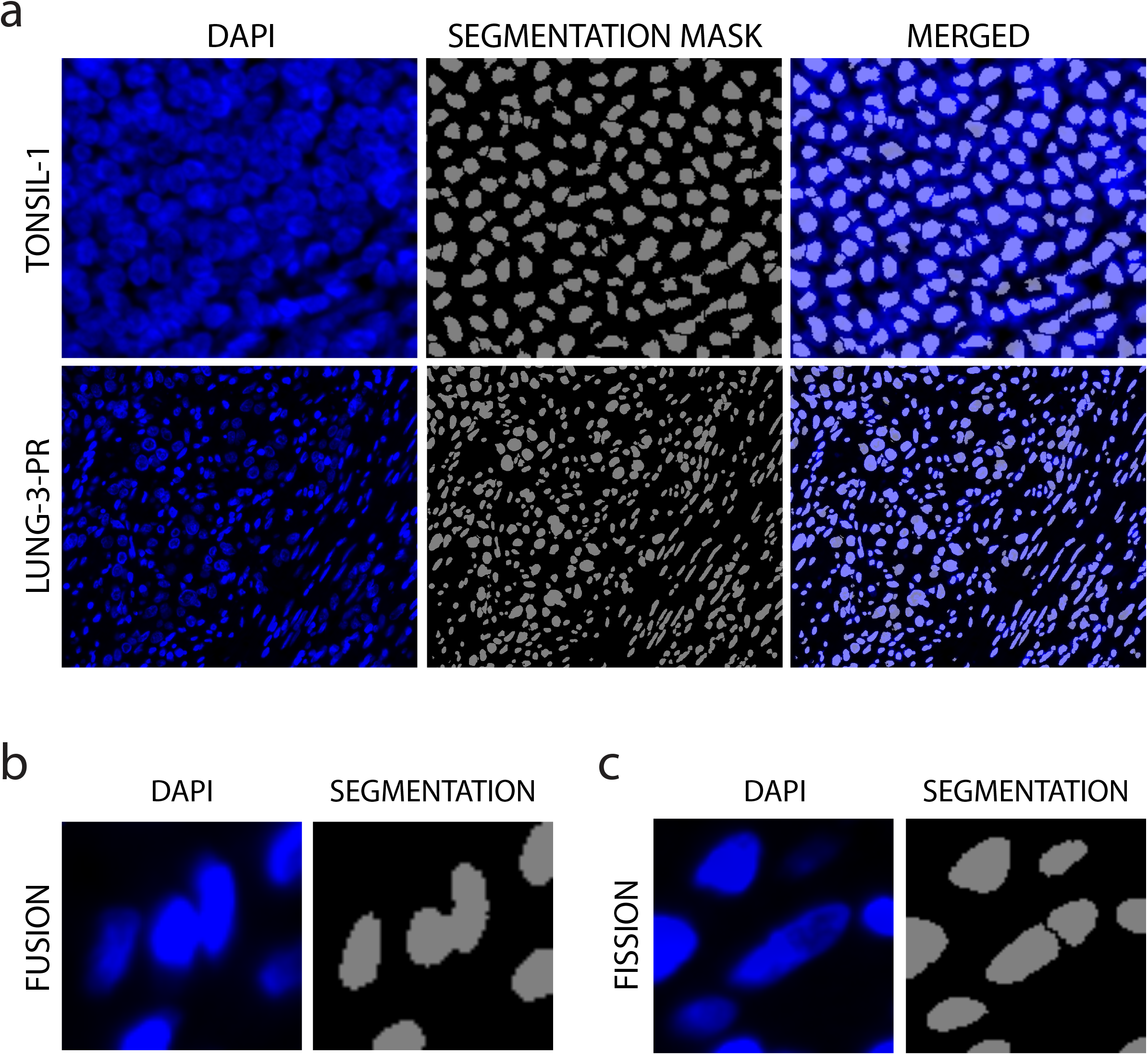
Assessment of segmentation. (a) Representative images of DAPI staining and corresponding segmentation mask in TONSIL-1 and LUNG-3-PR. (b) Examples of fusion (under-segmentation) and (c) fission/splitting (over-segmentation).

### Single-Cell Feature Extraction

To assess the integrity of the single-cell features extracted from the images, we applied an unsupervised, k-means clustering method to the data from the three lung cancer resection samples and the reactive tonsil sample. This yielded four cardinal cell types (clusters) using three lineage markers (Figure 5a). For each sample, the cells clustered into an epithelial group marked by keratin expression, a stromal group marked by αSMA expression, and a lymphocyte group marked by CD45 expression. A fourth group was marked by low expression of all three markers. We then isolated the cells in the lymphocyte group and further clustered them using other lymphocyte markers (Figure 5b-c). The clustering revealed similar immune cell populations to those observed by visual review of the images and as quantified using other computational methods in Du, Lin, Rashid *et al*., 2019^23^. Using alternative segmentation, feature extraction, and computational approaches, we retained reproducible immune cell populations, giving us confidence in the robustness of this dataset. However, it is highly probable that image segmentation can be further improved with the development of new algorithms.

**Figure 5.**
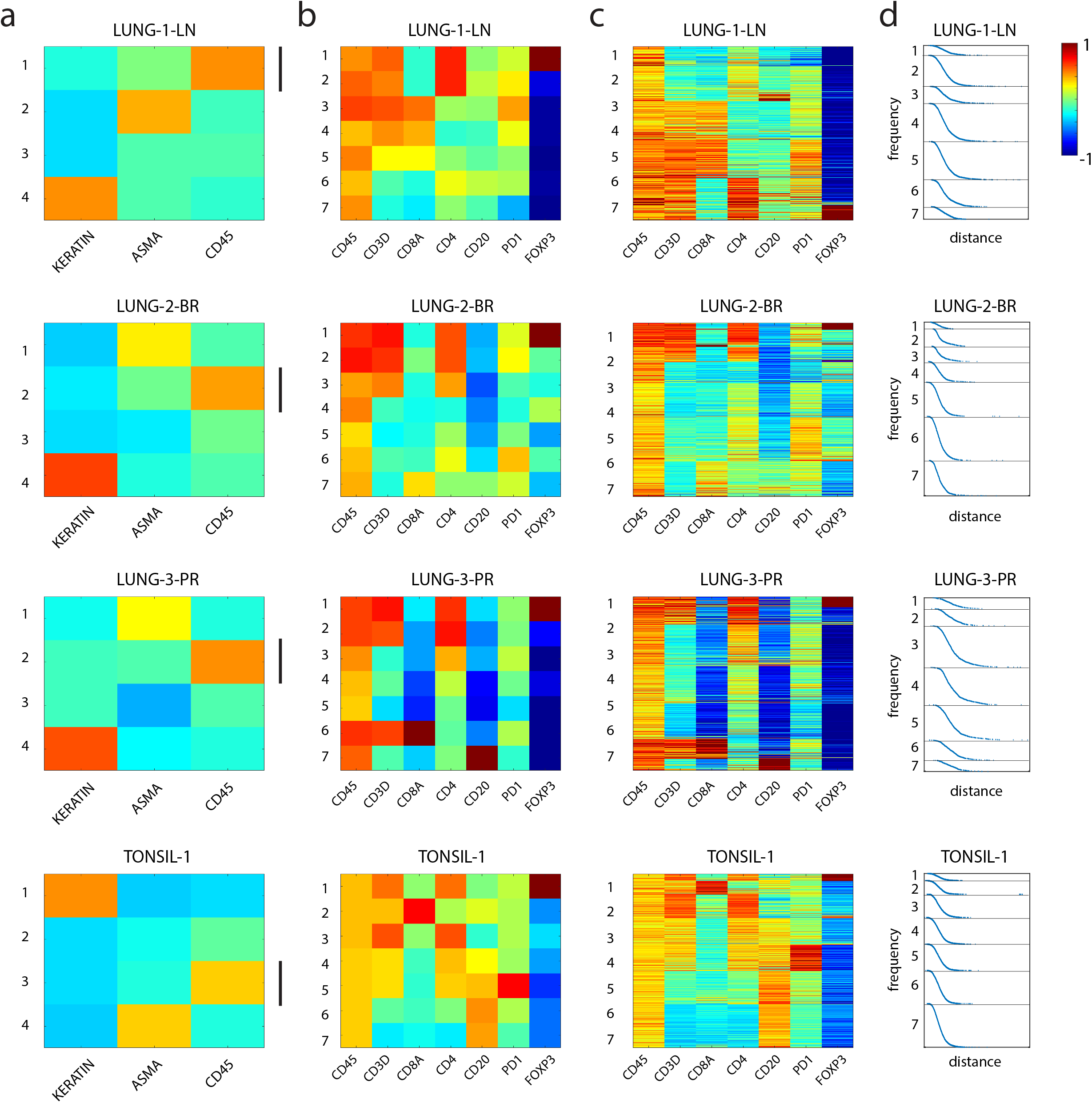
Heatmaps of cell populations from lung cancer and tonsil tissues using k-means clustering demonstrates distinct cell immune populations with expected patterns of biomarker expression. (a) Heatmap of the expression of Keratin, αSMA, and CD45 in all cells that were collected from LUNG-1-LN, LUNG-2-BR, LUNG-3-PR, and TONSIL-1 using k-means clustering. Each row is a cluster. The black vertical lines indicate the lymphocyte cluster with high CD45 expression from each tissue. (b. and c.) Heatmaps showing the expression of seven lymphocyte markers (CD45, CD3D, CD8A, CD4, CD20, PD1, FOXP3) from the cells within the CD45 high cluster from panel (a). (b) Each row represents a cluster or (c) each row represents protein marker expression data from a single cell. Note that data was log transformed and normalized between −1 to 1 as indicated by the color bar. (d) Plot showing fit of each cell within the cluster, with x-axis denoting the Euclidean distance from the centroid of the cluster and y-axis denoting the frequency of cells.

## USAGE NOTES

More information on the t-CyCIF method used to generate this data can be found at: www.cycif.org and a detailed protocol can be found in Lin *et al*., 2018^9^ and Du, Lin, Rashid *et al*., 2019^23^.

A narrative of the dataset is available for interactive web-browsing here: https://www.cycif.org/featured-paper/du-and-lin-2019/figures/

## ACKNOWLEDGEMENTS

This work was funded by NIH grants U54-CA225088 and U2C-CA233262 to P.K.S. and S.S., by U2C-CA233280 to P.K.S., and by the Ludwig Center at Harvard. The Dana-Farber/Harvard Cancer Center is supported in part by an NCI Cancer Center Support Grant P30-CA06516. D.S. was supported by the BioEntrepreneur-Fellowship of the University of Zurich (BIOEF-17-001) and an Early Postdoc Mobility fellowship (P2ZHP3_181475). G.G. was supported by T32-HL007627.

## AUTHOR CONTRIBUTIONS

R.R., G.G., Y.A.C., J.R.L., Z.D., Z.M., C.Y., D.S., J.M., and A.S. contributed to data collection, processing, and analysis. R.R., P.K.S., and S.S. wrote the manuscript. S.S. and P.K.S. supervised the project.

---

**Online-only Table 1:** Segmentation Accuracy

|  | LUNG-1-LN |  | LUNG-2-BR |  | LUNG-3-PR |  | TONSIL-1 |  |
| --- | --- | --- | --- | --- | --- | --- | --- | --- |
|  | Mean | SD | Mean | SD | Mean | SD | Mean | SD |
| **True Positive** | 0.872 | 0.013 | 0.89 | 0.017 | 0.899 | 0.013 | 0.863 | 0.012 |
| **Fusion** | 0.086 | 0.027 | 0.074 | 0.017 | 0.053 | 0.019 | 0.038 | 0.020 |
| **Fission** | 0.042 | 0.017 | 0.036 | 0.003 | 0.047 | 0.008 | 0.099 | 0.025 |

